# The macrophage reprogramming ability of antifolates reveals soluble CD14 as a potential biomarker for methotrexate response in rheumatoid arthritis

**DOI:** 10.1101/2021.05.11.443581

**Authors:** Sara Fuentelsaz-Romero, Celia Barrio Alonso, Raquel García Campos, Mónica Torres Torresano, Ittai Muller, Ana Triguero-Martínez, Laura Nuño, Alejandro Villalba, Rosario García-Vicuña, Gerrit Jansen, María-Eugenia Miranda-Carús, Isidoro González-Álvaro, Amaya Puig-Kröger

**Author notes:** Corresponding author: Dr. Amaya Puig-Kröger, Unidad de Inmunometabolismo e Inflamación, Instituto de Investigación Sanitaria Gregorio Marañón, Hospital General Universitario Gregorio Marañón, Madrid, Spain.

## Abstract

The physio-pathological relevance of the one-carbon metabolism (OCM) is illustrated by the chemotherapeutic and anti-inflammatory effects of the antifolates Pemetrexed (PMX) and Methotrexate (MTX) in cancer and rheumatoid arthritis (RA). We report that OCM determines the functional and gene expression profile of human macrophages. PMX induces the acquisition of a p53-dependent proinflammatory gene signature in human monocyte-derived macrophages (GM-MØ). Indeed, OCM blockade reprograms GM-MØ towards a state of LPS-tolerance at the signaling and functional levels, an effect abolished by folinic acid. Importantly, OCM blockade led to reduced expression of membrane-bound and soluble CD14 (sCD14), whose exogenous addition restores LPS sensitivity. The therapeutic relevance of these results was confirmed in early RA patients, as MTX-responder RA patients exhibit lower sCD14 serum levels, with baseline sCD14 levels predicting MTX response. Our results indicate that OCM is a metabolic circuit that critically mediates the acquisition of innate immune tolerance, and positions sCD14 as a valuable tool to predict MTX-response in RA patients.

## INTRODUCTION

Folates are one-carbon donors in biosynthetic pathways like *de novo* synthesis of purines and thymidylate, amino acid metabolism, and DNA methylation (1, 2). One-carbon metabolism (OCM) dispenses carbon atoms between various acceptor molecules required for biosynthesis and contributes to energy balance by providing ATP and NADPH (3). Accordingly, antifolates [including Methotrexate (MTX) and Pemetrexed (PMX)] are potent inhibitors of folate-dependent enzymes engaged in nucleotide synthesis, and inhibit DNA replication (2). PMX is a therapeutically relevant antifolate used in combination with cisplatin or carboplatin as the standard first-line treatment of malignant pleural mesothelioma (4, 5), advanced non-squamous non-small cell lung cancer (NSCLC) (6) and in maintenance therapy for NSCLC. PMX is transported into the cells via the Reduced Folate Carrier (RFC, *SLC19A1*) and, unlike MTX, is an excellent substrate for the proton-coupled folate transporter (PCFT, *SLC46A1*) (7). PMX is also an efficient substrate for the enzyme folylpoly-γ glutamate synthetase (FPGS) that catalyzes the addition of multiple glutamyl residues to the γ-carboxyl on the terminal glutamate of both folates and antifolates upon entry into the cell (8). PMX polyglutamylation enhances both its intracellular retention and its inhibitory potency towards thymidylate synthase (TS) and glycineamide ribonucleotide formyltransferase (GARFT) (8). However, whereas weekly administered MTX is the main starting therapy and the anchor drug for the treatment of rheumatoid arthritis (RA) (9), PMX is not approved for RA treatment, in spite of the fact that it reduces inflammatory cell infiltration in joints, limits cartilage-bone destruction and lowers serum TNFα and IL-17 in a rat model of collagen-induced arthritis (10).

Macrophages exhibit innate immune memory whereby they are durably primed by certain stimuli for enhanced or diminished responses to secondary stimuli (training/tolerance) (11–14). Macrophages are abundant in the synovium of RA joints, where they contribute to pathogenesis through the production of proinflammatory mediators (15–17). We have previously shown that macrophages from human RA joints exhibit a transcriptomic and phenotypic proinflammatory polarization profile that resembles that of GM-CSF-differentiated macrophages (GM-MØ) (18), and that MTX conditions macrophages towards the acquisition of a state of tolerance that renders them less responsive to TLR ligands, TNFα and RA synovial fluid (19, 20).

Low-dose MTX is safe, inexpensive and effective, but 30-40% of RA patients discontinue MTX treatment due to intolerance, inefficacy, or loss of clinical responsiveness at later time points (MTX resistance) (9, 21). Given these therapeutic limitations of MTX, and considering the widespread use of PMX in hyper-proliferative pathologies, we have now explored the phenotypic and functional effects of PMX on human macrophages. Our results indicate that PMX impairs TLR4 signaling and cytokine production in macrophages, and that these effects are dependent on the loss of membrane and soluble CD14 expression. The pathological significance of these findings has been confirmed in MTX-treated RA patients, which exhibit significantly reduced sCD14 levels. Altogether, our results indicate that PMX functionally reprograms human macrophages, lending support to its use as a potential replacement therapy for MTX, and demonstrate that innate tolerance can be induced in human macrophages through blockade of the one-carbon metabolism.

## METHODS

### Cell culture

Human peripheral blood mononuclear cells (PBMC) were isolated from buffy coats from normal donors over a Lymphoprep (Nycomed Pharma) gradient. Monocytes were purified from PBMC by magnetic cell sorting using CD14 microbeads (Miltenyi Biotech). Monocytes were cultured at 0.5 × 10^6^ cells/ml for 7 days in RPMI 1640 (standard RPMI, which contains 1 mg/L folic acid) supplemented with 10% fetal calf serum, at 37°C in a humidified atmosphere with 5% CO_2_, and containing GM-CSF (1000 U/ml) to generate GM-CSF-polarized macrophages (GM-MØ) or M-CSF (10 ng/ml) to generate M-CSF-polarized macrophages (M-MØ). GM-CSF or M-CSF was added every two days. Low-dose PMX (50 nM), MTX (50 nM) and/or folinic acid (FA, 5-formyl tetrahydrofolate, 5 μM), pifitrin-α cyclic (25-50 M) was added once on monocytes together with GM-CSF or M-CSF. When indicated, LPS (10 ng/ml, 0111:B4 strain, sLPS that exclusively binds TLR4) or LTA (5 μg/ml) were added at the indicated time points onto 7-day fully differentiated macrophages. sCD14 (20-200 ng/ml, Biolegend) was added on macrophages for intracellular signaling measurements (15 min) and IL-6 detection (3h).

### Rheumatoid Arthritis patients

#### Discovery cohort

Peripheral blood was obtained from 10 early rheumatoid arthritis patients who fulfilled 2010 American College of Rheumatology (ACR) revised criteria (22), with a disease duration of less than 6 months and who had never received disease-modifying drugs or corticosteroids. Plasma concentrations of sCD14 were determined by ELISA at the basal visit and 6 months after initiating treatment with weekly low-dose methotrexate (15-25 mg/week), none of the patients were taking prednisone at the time of the first or second determination. All of the included patients had a good clinical response to MTX (9 had achieved remission as determined by a DAS28<2.6 and the remaining one patient had persistent low activity with a ΔDAS28>1.2) and, accordingly, had discontinued prednisone at least one month prior to the 6 month sCD14 determination. Patients who required continued treatment with prednisone were not included in order to eliminate the possible interference of this immunomodulatory drug with the results. Disease activity was assessed using the DAS28 score (23). In sCD14 ELISA, interference of rheumatoid factor was ruled out after confirming that serial dilutions of plasma yielded identical sCD14 levels after adjusting for the diluting factor (24). The study was approved by the Hospital La Paz-IdiPAZ Ethics Committee, and all subjects provided written informed consent according to the Declaration of Helsinki.

#### Validation cohort

For sCD14 determination in serum, early arthritis patients in MTX monotherapy were recruited from the Princesa Early Arthritis Register Longitudinal (PEARL) study. The PEARL study comprises patients with one or more swollen joint and symptoms with ≤ 1 year of evolution. The register protocol includes 4 visits during a 2-year follow-up (0, 6, 12 and 24 months). Socio-demographic, clinical, therapeutic and laboratory data are recorded and included in an electronic database. Biological samples are collected at each visit and stored at −80°C in the Instituto de Investigación Sanitaria La Princesa (IIS-IP) Biobank for translational research. More detailed description of PEARL protocol has been previously published (25). None of the selected patients received corticosteroids prior to baseline visit, neither during the first 6 months of follow-up. Response was considered if at 6 months visit disease activity level decreased to remission or low disease activity provided that swollen joint count must have decreased, as well as must have decreased 3 out of the four following variables: tender joint count, erythosedimentation rate, C-reactive protein and global disease activity by patient. Non responders were considered if the disease activity level worsened from baseline to 6 months visit. The study was approved by the Hospital La Princesa Ethics Committee (PI-518, 28^th^ March 2011), and all subjects provided written informed consent according to the Declaration of Helsinki. Clinical and demographical data for discovery (Hospital La Paz) and validation (Hospital La Princesa) cohorts are:

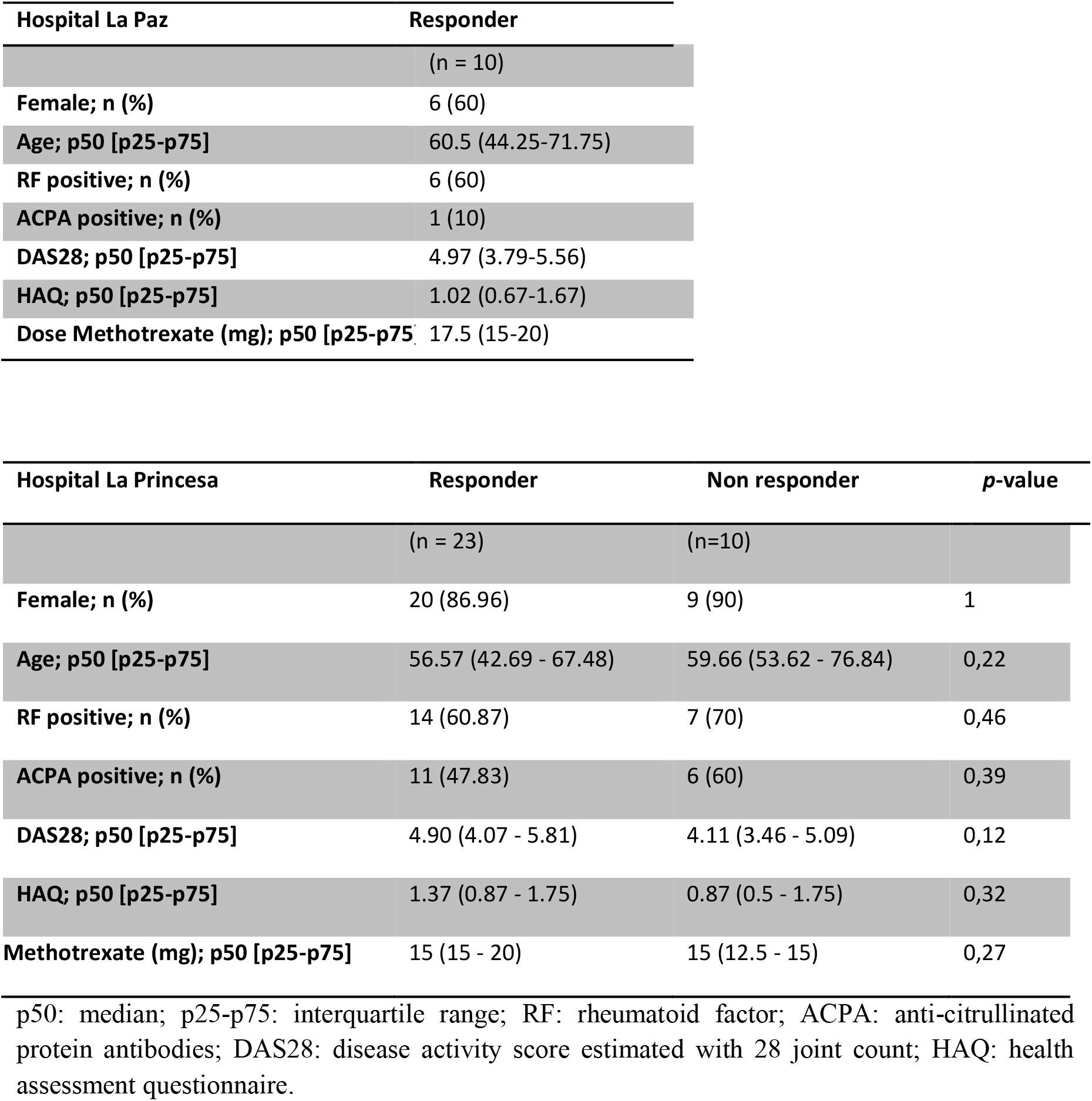

### RNAseq and GSEA

Total RNA was isolated from three independent preparations and processed at BGI (https://www.bgitechsolutions.com), where library preparation, fragmentation and sequencing were performed using the BGISEQ-500 platform. An average of 5.41 Gb bases were generated per sample and, after filtering, clean reads were mapped to the reference (UCSC Genome assembly hg38) using Bowtie2 (average mapping ratio 93.41%) (26). Gene expression levels were calculated by using the RSEM software package (27). Differential gene expression was assessed by using DEseq2 algorithms using the parameters Fold change>2 and adjusted *p* value <0.05. For gene set enrichment analysis (GSEA) (http://software.broadinstitute.org.gsea/index.jsp), the gene sets available at the website (28), as well as the “p53_Target_Gene” gene set, that contains the top 98 genes identified as p53 targets, were used (29). The data discussed in this publication have been deposited in NCBI’s Gene Expression Omnibus (30) and are accessible through GEO Series accession number GSE159349 and GSE159380.

### Quantitative real time RT-PCR

Total RNA was retrotranscribed and cDNA was quantified using the Universal Human Probe Roche library (Roche Diagnostics). Quantitative real-time PCR (qRT-PCR) was performed on a LightCycler^®^ 480 (Roche Diagnostics). Assays were made in triplicates and results normalized according to the expression levels of TBP. Results were obtained using the CT method for quantitation. The oligonucleotides used to quantify mRNA transcripts were (5′-3′): CD14s gttcggaagacttatcgaccat; CD14as acaaggttctggcgtggt; IL1βs ctgtcctgcgtgttgaaaga; IL1βas ttgggtaatttttgggatctaca; IL6s gatgagtacaaaagtcctgatcca; IL6as ctgcagccactggttctgt; TBPs cggctgtttaacttcgcttc; TBPas cacacgccaagaaacagtga.

### RNA interference

Two different siRNA for thymidylate synthase (*TYMS* silencer select s14538, s14539) and one control siRNA (silencer select Negative Control #1) (Ambion, Life Technologies) were used at 100 nM. GM-MØ were transfected with HiPerFect transfection reagent (QIAGEN), treated with PMX, and 48h later tested for CD14 mRNA levels by qRT-PCR.

### ELISA

Supernatants from GM-MØ were tested for the presence of IL-6, TNFα and CXCL10 (Biolegend), IFNβ (R&D Systems). For the detection of sCD14 in supernatants, serum and plasma of RA patients and healthy controls the Human CD14 Quantikine ELISA Kit (R&D Systems) was used.

### Western-blot

Cell lysates were obtained in RIPA buffer containing 1mM PIC (Protease Inhibitor Cocktail, SIGMA), 10 mM NaF, 1 mM Na_3_VO_4_ and 0.5 mM DTT. 10 micrograms cell lysate was subjected to SDS-PAGE and transferred onto an Immobilon polyvinylidene difluoride membrane (Millipore). Protein detection was carried out using mouse monoclonal Ab against IκBα (#4814; Cell Signaling), rabbit polyclonal against p-p38, pJNK and pERK (#9910; Cell Signaling), p-IRF3 (#4947; Cell Signaling) and p-STAT1 (#7649; Cell Signaling). Protein loading was normalized using an antibody against GAPDH (sc-32233, Santa Cruz Biotechnology) or against human Vinculin (#V4505; Sigma-Aldrich).

### Flow cytometry

Phenotypic analysis was carried out by immunofluorescence using standard procedures. Mouse monoclonal antibodies used for cell-surface staining included FITC-labeled anti-CD14 (BD) and PE-labeled anti-TLR4 (Biolegend). Isotype-matched labeled antibodies were included as negative controls.

### Statistical analysis

For the analysis of clinical data Stata 14.0 for Windows (Stata Corp LP, College Station, TX, USA) was used. For the discovery cohort a nonparametric Wilcoxon test was used to assess the statistical significance of differences between pre- and post-treatment RA plasma sCD14. For the validation cohort, most quantitative variables followed a non-normal distribution, so they were represented as median and interquartile range (IQR) and the Mann Whitney or Kruskal– Wallis tests were used to analyse significant differences. Qualitative variables were described using a calculation of the proportions and the Fisher’s exact test was used to compare categorical variables. To assess the ability of either baseline CD14 (sCD14) or the variation in sCD14 between baseline and six months follow-up visits (ΔsCD14) to discriminate between MTX-responder and MTX-non responder patients we generated Receiver Operating Characteristic (ROC) curves through the command roctab and the option graph. Each cut-off point was selected on the basis of the best trade-off values between sensitivity, specificity, cases correctly classified and positive (LR+) and negative (LR−) likelihood ratios reported using the command roctab with the option detail. Using these cut-off values new variables were generated to define patients with high or low baseline sCD14 and those with relevant sCD14 decrease. Then, we estimated the odds ratios and their 95% confidence interval with the command cs of Stata, with the option or. This command also provides the statistical significance with the Fisher exact test.

To estimate with a better precision the capability of sCD14 to discriminate between MTX-responder and non-responder patients, we performed a multivariable logistic regression analysis in which the dependent variable was response to MTX and as independent variables we included gender, age and baseline DAS28, that are well known predictors of treatment response in RA. Then, we consecutively included sCD14 or ΔsCD14 in the model to determine their respective OR adjusted by age, gender and baseline DAS28. A parametric Student’s t test was used to assess statistical significance between treatments and a *p* value < 0.05 was considered significant (^*^, *p*<0.05; ^**^, *p*<0.01, ^***^, *p*<0.001).

## RESULTS

### Pemetrexed induces a proinflammatory profile in GM-CSF-primed macrophages

We have previously described that low-dose MTX modifies the transcriptome of proinflammatory thymidylate synthase (TS)^+^ GM-CSF-primed macrophages (GM-MØ) (19) (GSE71253). Since PMX also influences the expression of MTX-regulated genes (*CCL20*, *LIF*, *MET* and *GDF15*) (19), we sought to determine the whole range of transcriptional changes induced by PMX in human macrophages. Comparison of the gene signatures of GM-CSF-primed monocyte-derived macrophages in the absence (GM-MØ) or presence (PMX-GM-MØ) of PMX (50 nM) (Figure 1A) revealed a large number of significant transcriptional differences (Figure 1B)., and that the gene profile of PMX-GM-MØ was enriched in terms like “HALLMARK_INFLAMMATORY_RESPONSE” and “TNFA_VIA_NF B_SIGNALING” GSEA gene sets (28) (Figure 1C, Supplementary Figure 1), supporting the notion that the continuous presence of PMX determines the acquisition of a more proinflammatory profile. Actually, PMX enhanced expression of NF B-target genes and MTX-regulated genes (Figure 1E), and, like MTX (19), potentiated the expression of p53 target genes (Figure 1D, F). Moreover, comparison of the gene signature of MTX-GM-MØ (GSE71253) and PMX-GM-MØ revealed a large overlap of the respective transcriptional effects of both antifolates (Figure 1G, H) (Supplementary Figure 1), thus emphasizing that, like MTX, PMX promotes the acquisition of a more proinflammatory and p53-dependent gene signature in GM-MØ.

**Figure 1.**
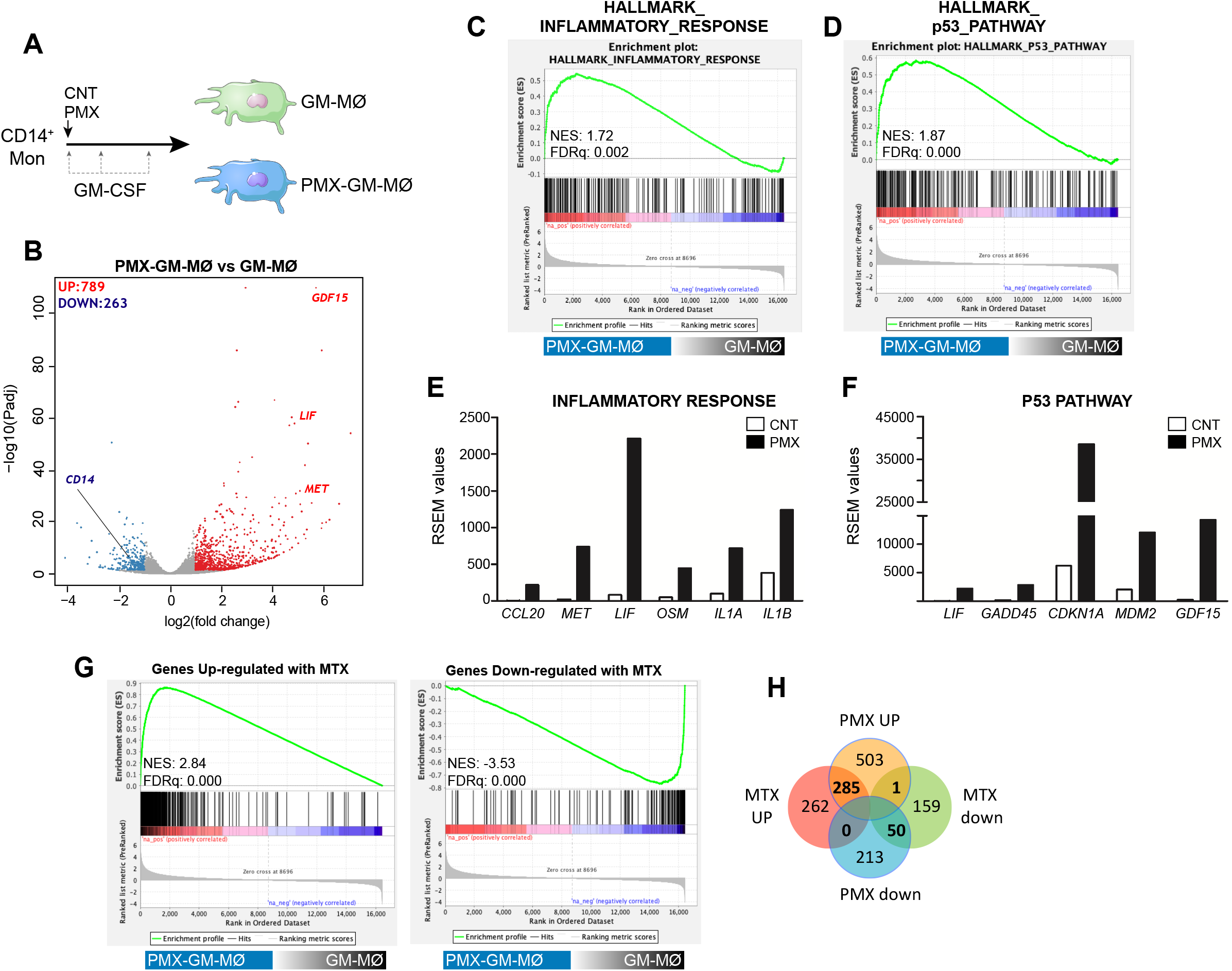
PMX promotes the acquisition of a proinflammatory gene profile in GM-MØ. **(A)** Schematic representation of the experiments. Monocytes were untreated or exposed to 50 nM PMX at the beginning of the 7-day macrophage differentiation process with GM-CSF and the mRNA levels were determined at day 7 on GM-MØ. **(B)** Volcano plot of RNAseq results, showing the PMX-induced gene expression changes in GM-MØ. The number of annotated genes whose expression is upregulated or downregulated in GM-MØ after 7d of PMX treatment (adjusted *p*< 0.05) is shown. **(C-D)** GSEA (http://software.broadinstitute.org/gsea/index.jsp) on the ranked list of genes obtained from the PMX-GM-MØ *versus* untreated GM-MØ limma analysis, using the Hallmarks v7 data set available at the web site. **(E-F)** RSEM expression values of the informative genes from the indicated GSEA enriched plot and found in the leading edge. **(G)** GSEA on the ranked list of genes obtained from the PMX-GM-MØ *versus* untreated GM-MØ limma analysis, using the genes significantly modulated by MTX in GM-MØ as data set. **(H)** Venn diagram comparing the genes differentially expressed by PMX in GM-MØ with the genes significantly altered by MTX in GM-MØ.

### PMX treatment promotes a transcriptional state of tolerance to LPS in human macrophages

Macrophage response to a given stimulus is dependent on extracellular cues that they have been previously exposed to (training/tolerance) (11–13). Thus, we next whether PMX promoted macrophage tolerance state at the transcriptional level. To that end, we determined the gene signature of LPS-stimulated (3h) GM-MØ differentiated in the presence of PMX, Folinic Acid (FA, a reduced folate with high-affinity for RFC (31)), or PMX+FA (Figure 2A). Compared to LPS-treated GM-MØ, LPS-treated PMX-GM-MØ exhibited a significantly (|log_2_FC| >1; adjp<0.05) elevated expression of 355 genes and diminished mRNA levels of 273 genes (Figure 2B). Supporting the specificity of PMX, none of these changes were observed in LPS-treated PMX+FA-GM-MØ, whose transcriptome was indistinguishable from that of control FA-GM-MØ (Figure 2B). Interestingly, GSEA revealed a very significant reduction of the genes within the GSEA “HALLMARK_INFLAMMATORY_RESPONSE” gene set (28) (Figure 2C) in LPS-treated PMX-GM-MØ, an enrichment that not seen in PMX+FA-GM-MØ (Figure 2C). Therefore, the presence of PMX during macrophage differentiation impairs the LPS-induced expression of inflammation-related genes, thus indicating that PMX promotes the acquisition of a tolerance state in macrophages. Conversely, expression of the “p53_Target_Genes” gene set was significantly and specifically enriched in LPS-treated PMX-GM-MØ (Figure 2C). Indeed, the role of p53 in the PMX-induced tolerance was demonstrated by using the p53 inhibitor pifitrin-α, that dose-dependently diminished the expression of PMX-responsiveness genes (*GDF15*, *LIF*, *CDKN1A* mRNA) in LPS-treated PMX GM-MØ (Figure 2D). Since the p53 target gene p21 determines the reparative and anti-inflammatory profile of macrophages (32), this result further supports the notion that PMX conditions macrophages towards a tolerance transcriptional state.

**Figure 2.**
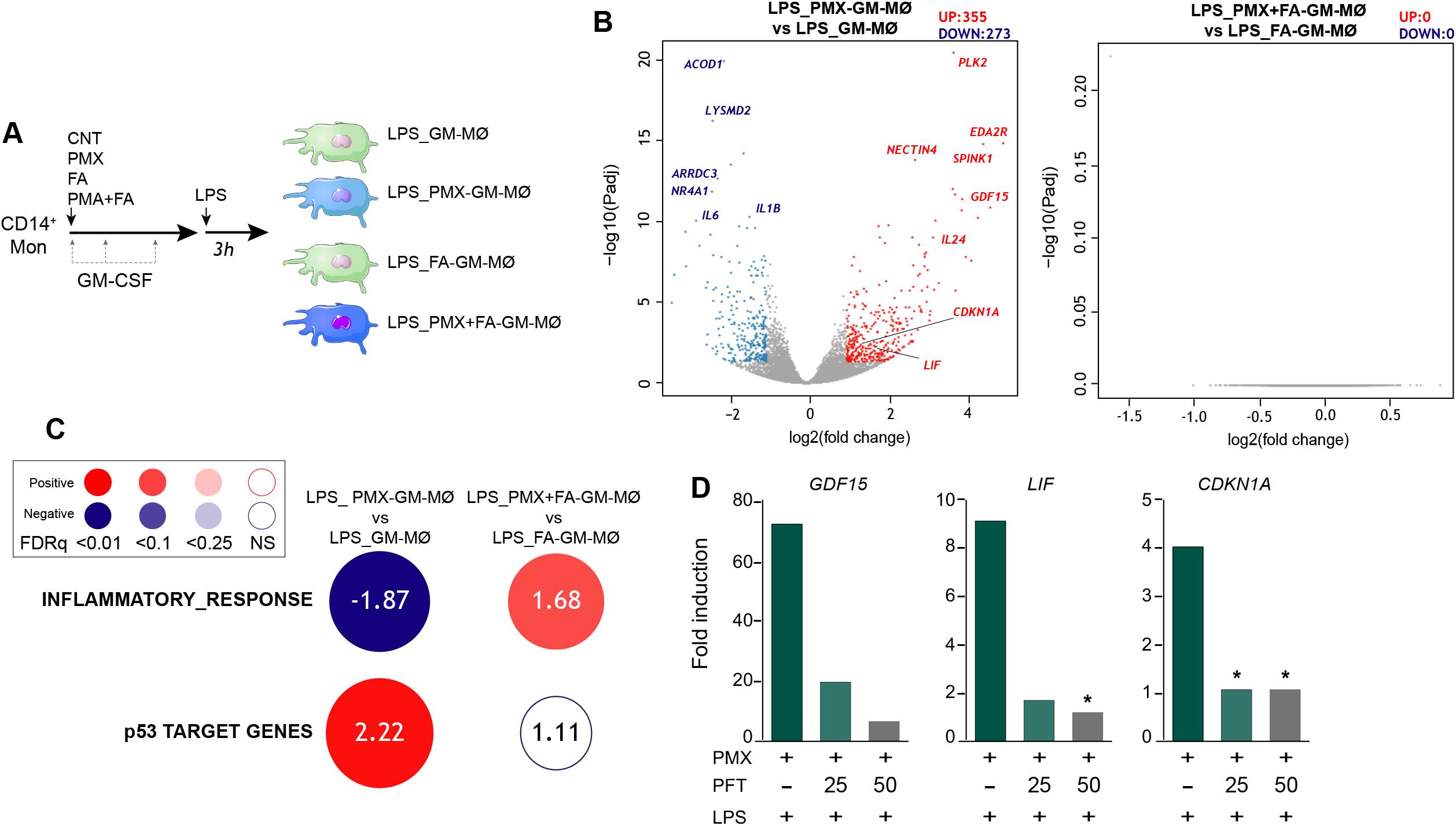
PMX promotes a transcriptional state of LPS-tolerance dependent on p53. **(A)** Experimental design. Monocytes were untreated (GM-MØ), exposed to 50 nM PMX (PMX-GM-MØ), 5 M folinic acid (FA-GM-MØ) or both (PMX+FA-GM-MØ) at the beginning of the 7-day macrophage differentiation process with GM-CSF and challenged with LPS (10 ng/ml) on day 7. Cells were assayed 3h post LPS-stimulation. **(B)** Volcano plot of RNAseq results, showing gene expression changes 3h post-LPS stimulation in PMX-treated GM-MØ (LPS_PMX-GM-MØ / LPS_GM-MØ, left), and PMX+FA-treated GM-MØ (LPS_PMX+FA-GM-MØ / LPS_FA-GM-MØ, right). The number of annotated genes whose expression is upregulated or downregulated 3h post-LPS stimulation in GM-MØ after 7d PMX, FA or PMX+FA treatment (adjusted *p*< 0.05) is shown. **(C)** Summary of GSEA with the indicated gene sets on the ranked comparison of the transcriptomes of LPS-treated PMX-GM-MØ *vs* LPS-treated GM-MØ, and LPS-treated PMX+FA-GM-MØ *vs* LPS-treated FA-GM-MØ. Circles area is proportional to the absolute value of the Normalized Enrichment Score (NES). The intensity of color increases with the enrichment of the gene signature (red, positive enrichment; blue, negative enrichment). False discovery rate (FDRq) is also indicated. The gene sets used were the Hallmarks v7 data set available at the web site and the “p53_Target_Gene” data set (29). **(D)** Gene expression determined by qRT-PCR on monocytes stimulated with PMX in the absence or presence of pifitrin- (PFT, 25-50 M) during the GM-CSF-dependent differentiation process and challenged with LPS for 3h. Mean ± SEM of 4 independent donors are shown (^*^*p*<0.05). Results are expressed as fold induction, which indicates the expression of each gene in PMX-exposed relative to control cells, and untreated or treated with PFT.

### PMX attenuates LPS-induced inflammatory cytokine production in human macrophages: involvement of the TLR4 signaling pathway

To evaluate the functional correlate of the PMX-induced transcriptional tolerance, we analyzed the ability of PMX to modify human macrophage responses upon TLR4 engagement. In agreement with the gene ontology analysis, LPS-treated (3h) PMX-GM-MØ displayed a considerably diminished *IL6* and *IL1B* mRNA expression when compared to LPS-treated GM-MØ (Figure 3A). Moreover, PMX-GM-MØ produced significantly lower levels of LPS-induced IL-6, TNFα and IL-10 after both short-term (3h) and long-term (24h) LPS exposure (Figures 3B-D). The specificity of the PMX effect was demonstrated by the absence of the inhibitory effect in PMX+FA-GM-MØ (Figure 3A, E), thus indicating that the effects of PMX are mediated through OCM blockade. Of note, PMX-GM-MØ also produced lower level of IL-6 after exposure to the TLR2 ligand lipoteichoic acid (LTA), thus indicating that PMX attenuates inflammatory cytokine expression in response to either TLR4 or TLR2 ligands. Altogether, these results indicate that PMX conditions macrophages for the acquisition of a tolerance state at the transcriptional and functional levels.

**Figure 3.**
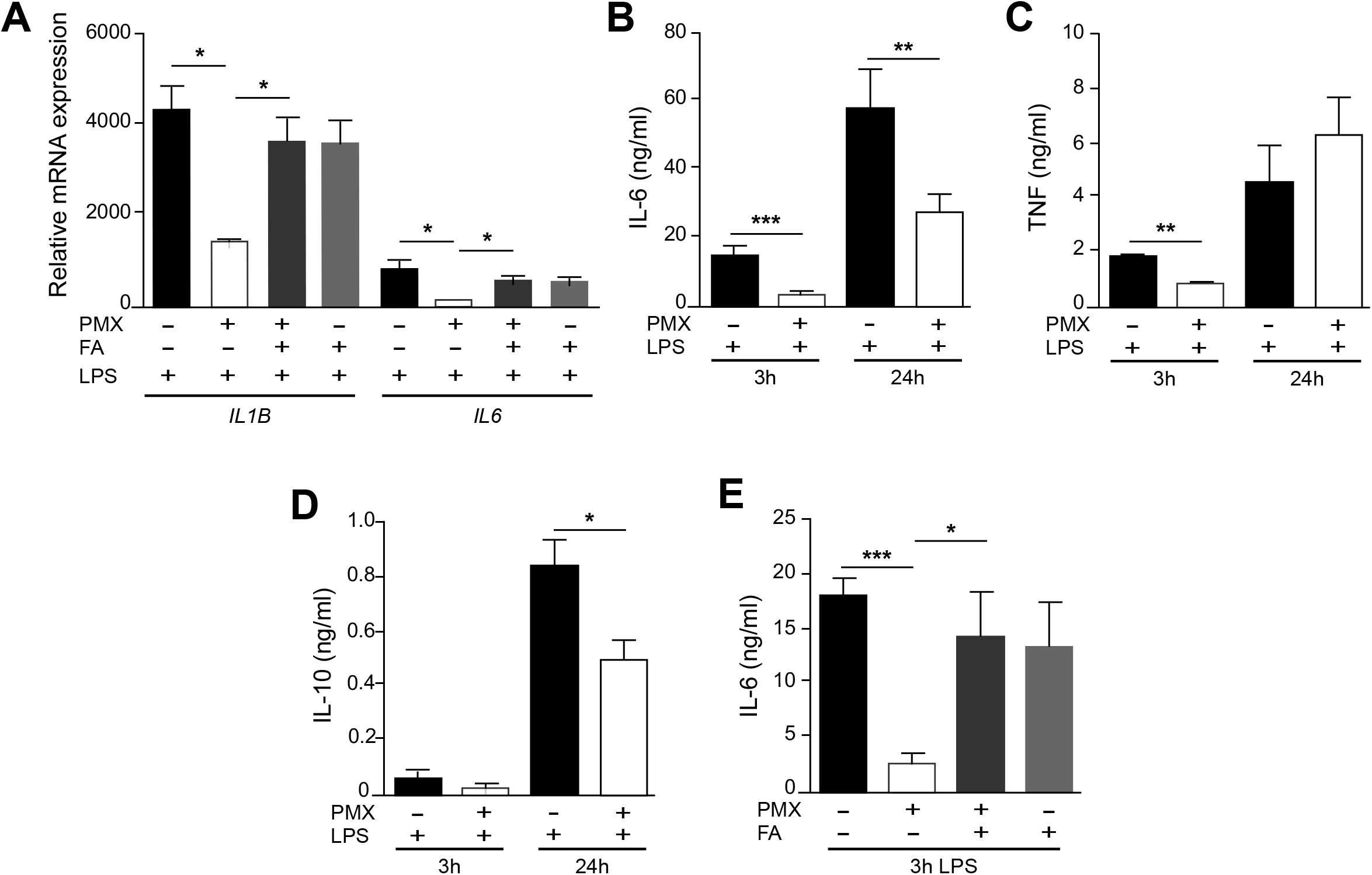
PMX alters TLR4 responsiveness in GM-CSF-primed macrophages. **(A)** Expression of *IL1B* and *IL6* by monocytes differentiated with GM-CSF in the absence or presence of PMX (50 nM), folinic acid (FA, 5 μM) or both and challenged with LPS for 3h, as determined by qRT-PCR. Mean ± SEM of 3 independent donors are shown (^*^*p*<0.05). Production of IL-6 **(B)**, TNFα **(C)** or IL10 **(D)** by monocytes differentiated with GM-CSF in the absence or presence of PMX and challenged with LPS for 3h and 24h, as determined by ELISA. Mean ± SEM of 7 independent donors are shown (^*^*p*<0.05, ^**^*p*<0.01, ^***^*p*<0.001). **(E)** Production of IL-6 by monocytes differentiated with GM-CSF in the absence or presence of PMX (50 nM), folinic acid (FA, 5 μM) or both and challenged with LPS for 3h, as determined by ELISA. Mean ± SEM of 5 independent donors are shown (^*^*p*<0.05, ^***^*p*<0.001).

Previous studies have demonstrated that tolerant-macrophages exhibit a lower level of MAPK and NFκB activation upon TLR stimulation (20, 33, 34). To determine whether PMX promotes “bonafide” innate tolerance in macrophages, we first determined the levels of MAPK and IκBα activation in LPS-treated PMX-GM-MØ. Compared to untreated PMX-GM-MØ, LPS-treated PMX-GM-MØ exhibited a reduced activation of p38 and JNK (Figure 4A). By contrast, and as described in other cellular systems (35), PMX-GM-MØ showed an increased and more sustained phosphorylation of ERK1/2 in response to LPS (Figure 4A). Besides, a lower level of IκBα degradation was detected in LPS-exposed PMX-GM-MØ (Figure 4B), a reduction abolished in the presence of folinic acid (PMX-FA-GM-MØ, Figure 4C). Further and compared to GM-MØ, LPS induced lower levels of p-IRF3, p-STAT1, and, IFNβ1 and CXCL10 secretion in PMX-GM-MØ (Figure 4D-G). Taken together, these results demonstrate that OCM mediates the impaired TLR4-initiated intracellular signaling and the establishment of the state of LPS tolerance promoted by PMX, and suggest that PMX exerts its effects primarily via the TRIF-type I IFN branch of the TLR4 intracellular signaling (36).

**Figure 4.**
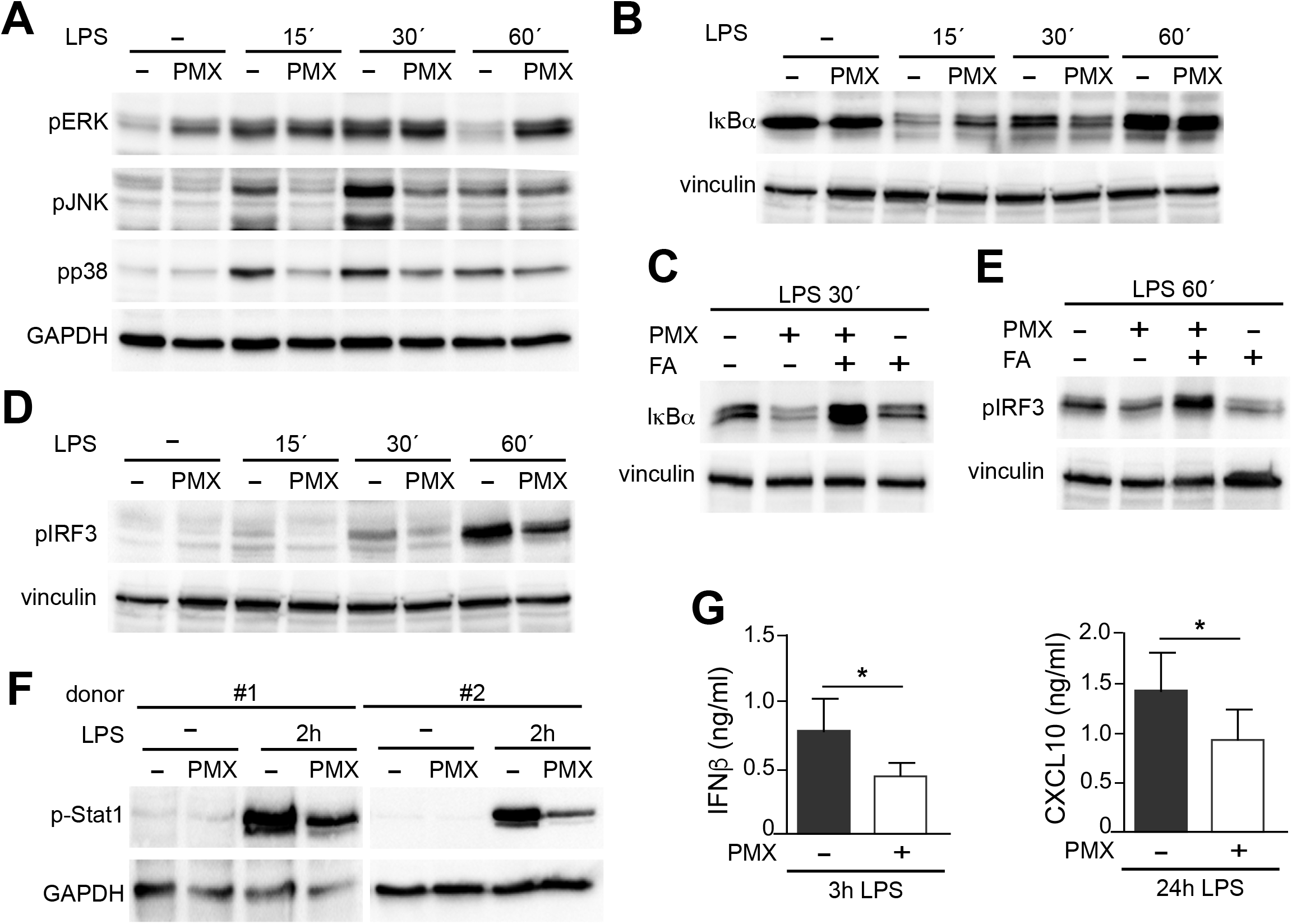
PMX alters TLR4 signaling in GM-CSF-primed macrophages. **(A-B)** Immunoblot analysis of pp38, pJNK and pERK **(A)** and IκBα **(B)** by monocytes differentiated with GM-CSF in the absence or presence of PMX for 7 days and challenged with LPS for the indicated time points. **(C)** Immunoblot analysis of I B by monocytes differentiated with GM-CSF in the absence or presence of PMX, PMX+FA or FA for 7 days and challenged with LPS for 30 minutes. **(D-E)** Immunoblot analysis of pIRF3 by monocytes differentiated with GM-CSF in the absence or presence of PMX **(D)**, folinic acid (FA) or both **(E)** for 7 days and challenged with LPS for the indicated time points. **(F)** Immunoblot analysis of pStat1 by two independent preparations of monocytes differentiated with GM-CSF in the absence or presence of PMX for 7 days and challenged with LPS for 2h. In A-F a representative experiment of 2-4 independent donors is shown. **(G)** Production of IFN and CXCL10 by monocytes differentiated with GM-CSF in the absence or presence of PMX challenged with LPS for 3h (IFN) or 24h (CXCL10), as determined by ELISA. Mean ± SEM of 7 independent donors are shown (^*^*p*<0.05).

### PMX and MTX regulate CD14 expression in GM-CSF-primed human macrophage

Next, we determined whether PMX affected the expression of negative regulators of TLR-induced cytokine production that contribute to LPS tolerance (33). Unlike MTX-induced tolerance, mostly mediated by increased *TNFAIP3* (A20) expression (33, 34, 37, 38), no difference in *TNFAIP3* expression was found between LPS-treated GM-MØ and LPS-treated PMX-GM-MØ (not shown). Thus, we next screened for genes coding for regulators of TLR-induced activation and whose expression significantly differs between GM-MØ and PMX-GM-MØ, and found that PMX-GM-MØ express significantly lower levels of *CD14* mRNA than GM-MØ (Figure 5A). After validation of this result in additional samples (Figure 5A), we confirmed that PMX-GM-MØ express a significantly lower level of cell surface CD14 (mCD14) than GM-MØ (Figure 5B-C). Specifically, the percentage of CD14-positive cells was 75% in GM-MØ and 45% in PMX-GM-MØ, whereas the mean fluorescence significantly varied from 46.5 to 36, respectively (not shown). Further, the soluble form of CD14 (sCD14) was also significantly lower in the supernatant of PMX-GM-MØ than in control GM-MØ cells (Figure 5D), and the PMX-induced reduction of mCD14 and sCD14 was completely reversed in the presence of folinic acid (Figure 5C, 5E). Of note, the PMX-triggered CD14 downregulation was significantly diminished after siRNA-mediated *TS* knockdown (Figure 5F), thus indicating that the PMX-triggered CD14 downregulation is dependent on TS expression. Moreover, knockdown of TS sufficed to diminish CD14 mRNA expression in GM-MØ. Of note, the antifolate MTX exhibited the same effect, as MTX-GM-MØ showed significantly lower expression of *CD14* mRNA, mCD14 and sCD14 than GM-MØ, an effect also prevented by folinic acid (Figure 5G-K). Therefore, OCM mediates the loss of *CD14* gene and protein expression in macrophages differentiated in the presence of antifolates (PMX-GM-MØ or MTX-GM-MØ). The physiological significance of this findings was further stressed by the fact that both antifolates only impaired CD14 expression in proinflammatory GM-MØ, whereas they did not modify CD14 expression in M-CSF-conditioned macrophages (M-MØ) (Figure 5G-H), which resemble anti-inflammatory tissue-resident and pro-tumoral TAM (18, 39, 40).

**Figure 5.**
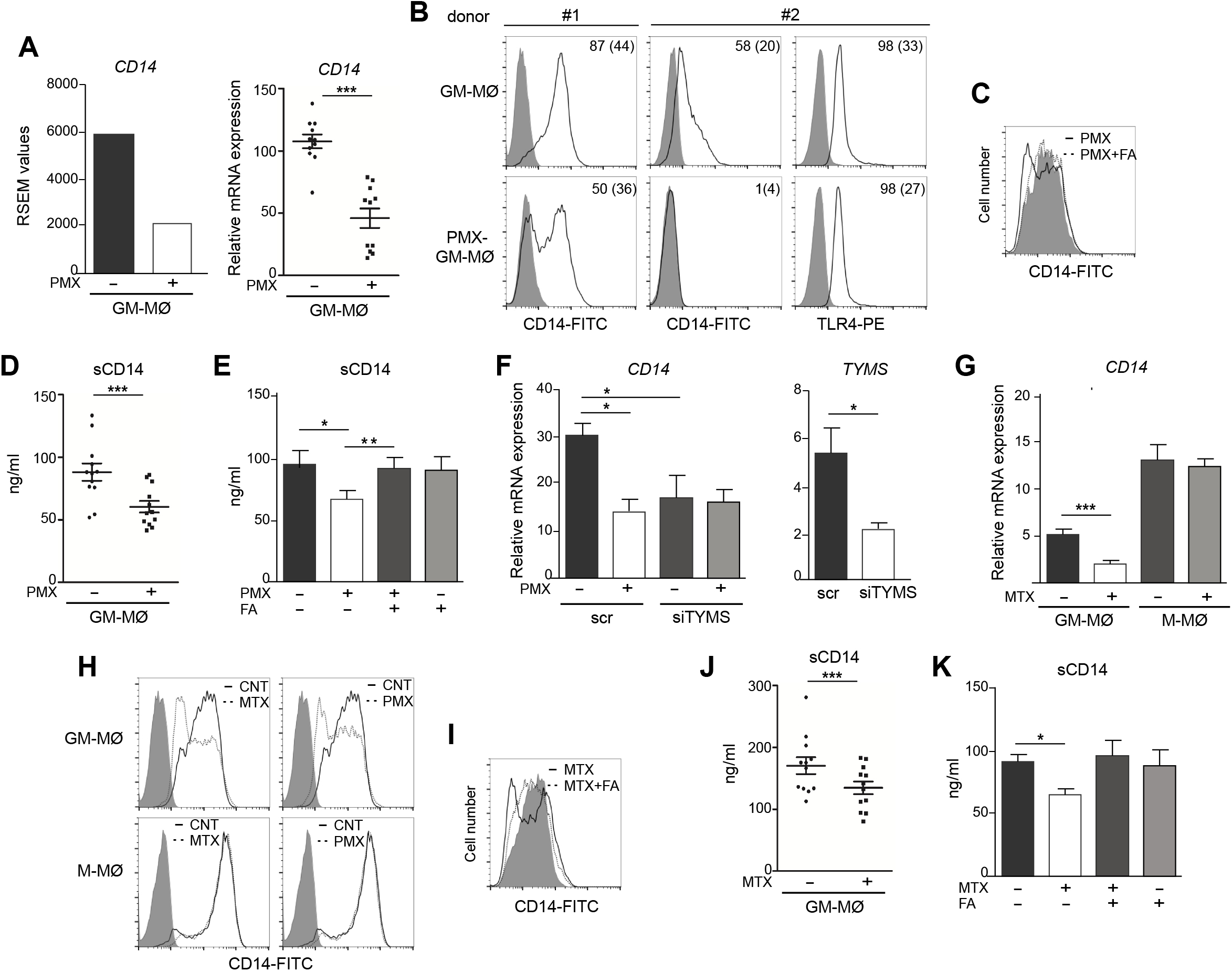
OCM blockade diminish CD14 expression on GM-CSF-primed macrophages. **(A)** *CD14* mRNA expression on monocytes stimulated with PMX during the GM-CSF-dependent differentiation process, as determined by RNAseq (adj*p*=10^−5^, left) and by qRT-PCR (right, n=11 independent donors, each symbol represent a single donor, ^***^*p*<0.001). **(B)** Cell surface expression of CD14 and TLR4 on monocytes differentiated with GM-CSF in the absence or in the presence of PMX, as determined by flow cytometry. The percentage of marker-positive cells and the mean fluorescence intensity (in parentheses) are indicated. The experiment was done on 8 independent donors, and 2 representative donors are shown. **(C)** Cell surface expression of CD14 on monocytes differentiated with GM-CSF in the absence (filled histograms) or in the presence of PMX (unfilled line histograms) or PMX+FA (dotted line histograms), as determined by flow cytometry. The experiment was done on 4 independent donors, and a representative is shown. **(D)** Soluble CD14 level in the supernatant of monocytes differentiated with GM-CSF in the absence or in the presence of PMX, as determined by ELISA, (n=12 independent donors, each symbol represent a single donor; ^***^*p*<0.001). **(E)** Soluble CD14 level in the supernatant of monocytes differentiated with GM-CSF in the absence or presence of PMX, folinic acid (FA) or both, as determined by ELISA (n=6 independent donors). **(F)** Left, *CD14* mRNA expression on GM-MØ transfected with control siRNA (scr) and siRNA for thymidylate synthase (siTYMS) and exposed to PMX for 48 hours, as determined by qRT-PCR (n=4 independent donors, ^*^*p*<0.05). Right, *TYMS* mRNA expression on GM-MØ transfected with control siRNA and siRNA for TYMS for 48 hours, as determined by qRT-PCR (n=4 independent donors, ^*^*p*<0.05). **(G)** *CD14* mRNA expression on monocytes stimulated with MTX (50 nM) during the GM-CSF or M-CSF-dependent differentiation process, as determined qRT-PCR (n=6 independent donors, ^***^*p*<0.001). **(H)** Cell surface expression of CD14 on monocytes differentiated with GM-CSF (GM-MØ) or M-CSF (GM-MØ) in the absence or in the presence of MTX or PMX, as determined by flow cytometry. **(I)** Cell surface expression of CD14 on monocytes differentiated with GM-CSF in the absence (filled histograms) or in the presence of MTX (unfilled line histograms) or MTX+FA (dotted line histograms), as determined by flow cytometry. For H-I, the experiment was done on four independent donors, and a representative is shown. **(J)** Soluble CD14 level in the supernatant of monocytes differentiated with GM-CSF in the absence or in the presence of MTX, as determined by ELISA, (n=15 independent donors, each symbol represent a single donor; ^***^*p*<0.001). **(K)** Soluble CD14 level in the supernatant of monocytes differentiated with GM-CSF in the absence or presence of MTX, folinic acid (FA) or both, as determined by ELISA (n= 4 independent donors, ^*^*p*<0.05).

### Exogenous sCD14 restores the impaired LPS-responsiveness of PMX or MTX-GM-MØ

Since the reduced expression of sCD14 and mCD14 correlated with diminished intracellular signaling and cytokine responses to LPS in PMX-GM-MØ, we next determined whether CD14 had a role in the PMX-induced macrophage tolerance state by assaying LPS-responsiveness of PMX-GM-MØ in the presence of increasing concentrations of exogenous sCD14. Addition of sCD14 increased the LPS-induced activation of p38 and JNK, as well as the LPS-induced degradation of IκBα in PMX-GM-MØ, but not in GM-MØ (Figure 6A). Noteworthy, the same results were found in MTX-GM-MØ (Figure 6B). Moreover, sCD14 dose-dependently increased the LPS-induced IL-6 production in PMX-GM-MØ, but not in LPS-treated GM-MØ (Figure 6C). Therefore, exogenous *s*CD14 partly restores the inhibitory effect of PMX on TLR4-initiated intracellular signaling and IL-6 production, and demonstrate that CD14 mediates the OCM-dependent pro-tolerant effect of PMX.

**Figure 6.**
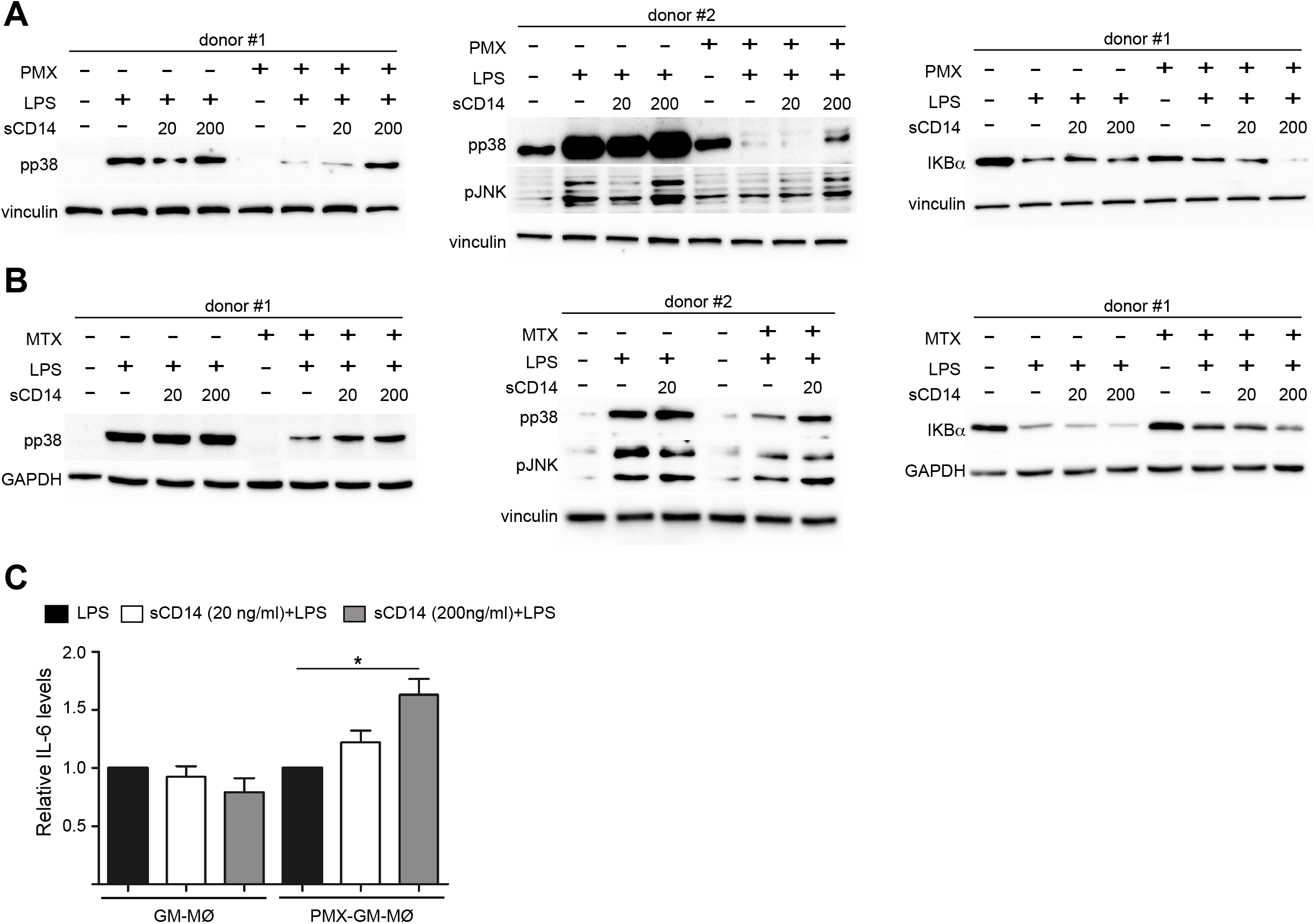
sCD14 restores p38 and JNK activation and IL-6 production in LPS-treated PMX-GM-MØ. Immunoblot analysis of pp38, pJNK and IκBα by GM-MØ and PMX-GM-MØ **(A)** or MTX-GM-MØ **(B)** challenged with LPS in the absence or presence of 20 ng/ml (20) or 200 ng/ml (200) of sCD14 for 15 min. (pp38, pJNK) and 30 min. (IκBα). The experiment was performed in 4 independent donors and 2 of them are shown. The signal in pp38 in donor #2 (A) is saturated for better detection of lane 8 (sCD14 200) in PMX-GM-MØ. **(C)** Production of IL-6 by GM-MØ or PMX-GM-MØ challenged with LPS in the absence or presence of 20 ng/ml or 200 ng/ml of sCD14 for 3h, as determined by ELISA. The experiment was performed in 4 independent donors and the relative production of IL-6 is shown.

### Soluble CD14 diminishes in early rheumatoid arthritis patients who are methotrexate-responders

Finally, we sought to determine the therapeutic relevance of the CD14 downregulation in GM-MØ generated in the presence of antifolates. Although PMX and MTX similarly diminished mCD14 and sCD14 in GM-MØ (Figure 5) only low-dose MTX is an anchor drug for RA treatment (9, 41). Therefore, we turned to RA patients treated with MTX to evaluate the clinical significance of the association between antifolate exposure and CD14 downregulation. Specifically, we determined sCD14 level in plasma from MTX-responder RA patients both at baseline and after 6 months of treatment with MTX. Remarkably, we found that sCD14 levels significantly decreased in patients who respond to MTX (Figure 7A). Indeed, sCD14 levels positively correlated with disease activity score DAS28 and CRP (Supplementary Figure 2). These results indicate that sCD14 expression is lower in MTX responder RA patients, suggesting that sCD14 might be a biomarker for MTX-response. To confirm these findings, we determined sCD14 concentration in a validation cohort of early arthritis patients at baseline and during 6 months of MTX monotherapy, including both MTX-responder and non-responder patients (see methods for definition). Interestingly, baseline sCD14 levels in MTX non-responder patients were similar to control healthy donors, whereas MTX-responder patients exhibited significantly higher sCD14 serum levels (Figure 7B). Moreover, sCD14 levels diminished in MTX-responder patients but not in MTX non-responders (Figure 7C). In order to determine whether baseline sCD14 and ΔsCD14 could be MTX response biomarkers, we performed ROC analysis. Although the area under curve (AUC) of ΔsCD14 was slightly higher than that of baseline sCD14 (Figure 7D), it did not reach statistical significance. The best cutoff to discriminate between MTX responder and non-responder patients was 2.460 ng/ml for baseline sCD14 (78% sensitivity and 70% specificity) and 188 ng/ml for ΔsCD14 (83% sensitivity and 70% specificity). The odd ratio (OR) for having high baseline sCD14 and being MTX-responder was 8.4 (p=0.0126), whereas OR for a decreased ΔsCD14 was 11.1 (p=0.0059) (Figure 7E). Furthermore, after adjustment by variables related to MTX-response such as age, gender and baseline DAS28, OR increased to 25.5 (for sCD14) and 40.35 (for ΔsCD14) (Supplementary Table 1). Altogether, these results indicate that determination of sCD14 levels could be a valuable tool to predict or evaluate MTX-response in RA patients.

**Figure 7.**
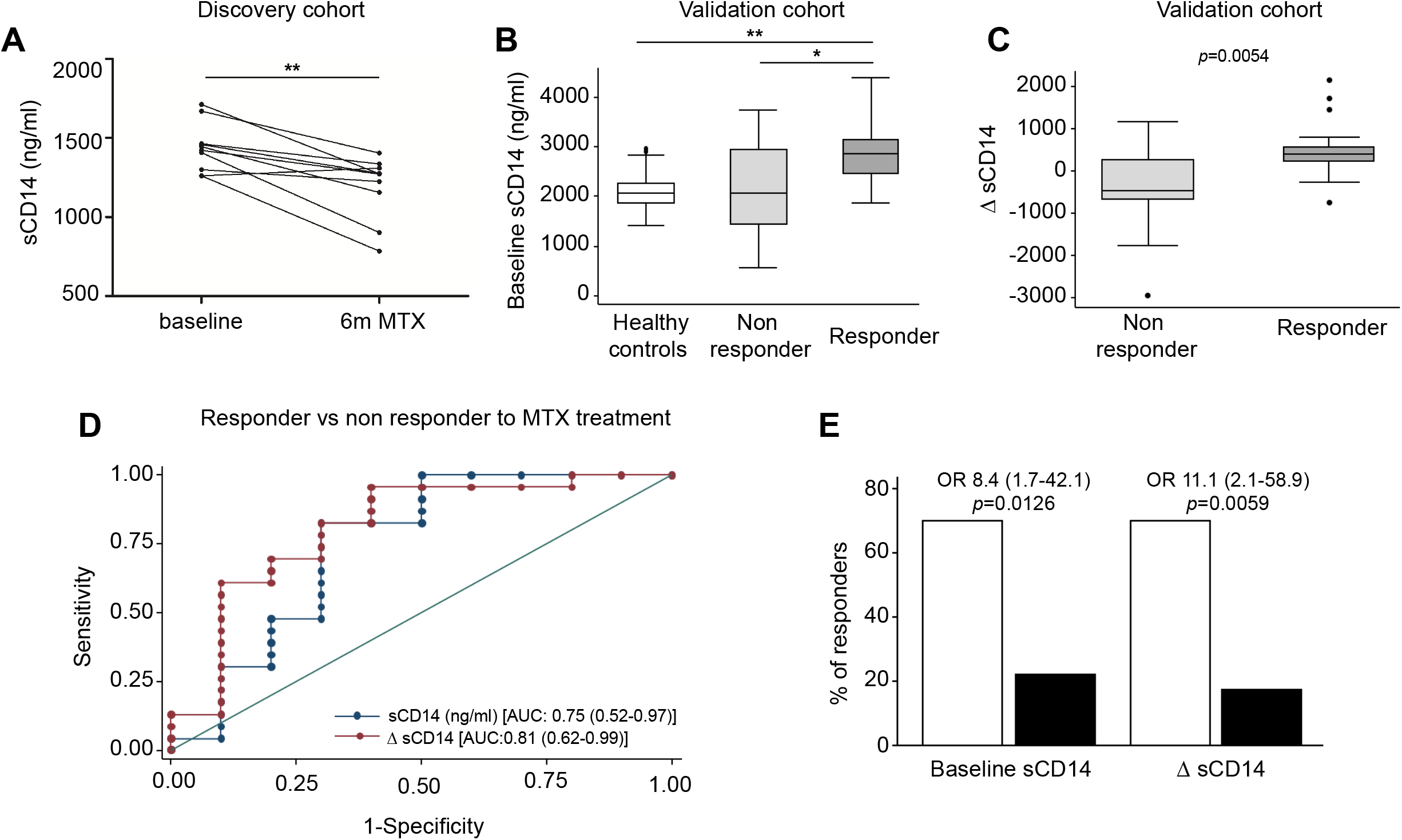
Serum level of sCD14 decreases in RA patients who are MTX-responders. **(A)** sCD14 level in plasma from early RA patients at baseline and 6 months after initiating MTX treatment (15-25 mg/week), as determined by ELISA (n=10, ^**^*p*<0.01, Wilcoxon test). **(B)** Baseline sCD14 in healthy donors (white box, n=40), MTX-responder (dark grey box, n=23) and MTX non-responder (soft grey box, n=10) RA patients. Statistical significance respect to MTX-responders was determined through Mann-Whitney test. **(C)** Variation in sCD14 between baseline and six months follow-up visits (ΔsCD14) in MTX-responder (dark grey box) and MTX-non responder (soft grey box) RA patients. Data in B-C are shown as interquartile range (p75 upper edge of box, p25 lower edge, p50 midline) as well as the p95 (line above box) and p5 (line below). **(D)** Receiver operating characteristic curves analysis to assess capacity of baseline sCD14 (blue line and dots) and ΔsCD14 (red line and dots) to discriminate between MTX-responder and MTX non-responder RA patients. **(E)** Percentage of MTX responders: comparison among patients with high (white bar) *versus* low (black bar) baseline sCD14 (left bars); comparison between patients with relevant decrease (white bar) *versus* non-relevant decrease plus increase (black bars) of sCD14 (right bars). Odds ratio and its confidence interval were estimated with the cs command of Stata 14.1 and the significance level with the Fisher exact test.

## DISCUSSION

One-carbon metabolism (OCM) is a complex network of biosynthetic pathways that includes *de novo* biosynthesis of purines and thymidylate, amino acid metabolism, and methylation reactions (1). In the present report, we describe the impact of OCM on the functional and gene expression profile of GM-CSF-primed human monocyte-derived macrophages (GM-MØ). We have found that OCM-blocker PMX induces the acquisition of a more proinflammatory and p53-dependent gene signature in human macrophages, an effect that resembles the transcriptomic changes triggered by MTX, another antifolate with OCM-blocking activity (Figure 1) (19). Moreover, we describe that blocking OCM reprograms GM-MØ towards a tolerance state, as PMX-GM-MØ exhibit a reduced response to TLR ligands. The specificity of PMX effects was demonstrated by the absence of the inhibitory effect in the presence of folinic acid, a reduced folate with high-affinity for RFC (31). Mechanistically, PMX reduces LPS-induced p38, JNK, IRF3 and STAT1 activation, IκBα degradation and inflammatory cytokine production in human macrophages. In line with these findings, PMX also diminished the expression of membrane-bound and soluble CD14, a co-receptor for TLR4 required for MyD88-dependent signaling at low concentrations of LPS, and for LPS-induced TLR4 internalization into endosomes and activation of TRIF-mediated signaling (42, 43). Accordingly, PMX-GM-MØ exhibit diminished MyD88-dependent and TRIF-mediated signaling, as well as reduced cytokine production (IL-6, IFNβ1), upon exposure to LPS. The relevance of sCD14 in the OCM-dependent pro-tolerant effect of PMX is supported by the ability of exogenous sCD14 to restore the LPS sensitivity of PMX-GM-MØ. Altogether, these results indicate that the global anti-inflammatory activity of PMX relies on its ability to induce a proinflammatory profile in GM-CSF-primed macrophages, making PMX-conditioned macrophages less responsive to secondary inflammatory stimuli. These results link OCM to innate immune tolerance (44), and demonstrate that antifolates promote a proinflammatory state in macrophages and, in parallel, trigger a loss of CD14 expression, all of which end up establishing a tolerant state and an impaired response to subsequent stimulation.

Cellular metabolism is a critical mediator of the reprogramming of myeloid cells that takes place during trained immunity (45). Increased aerobic glycolysis is a hallmark of β-glucan or BCG-induced trained immunity in monocytes (46, 47). Regarding innate tolerance, the metabolite itaconate inhibits LPS-mediated IκB induction and induces tolerance in human monocytes (48). The lipid and amino acid metabolisms are also important for the induction of trained immunity (45), as metabolites of the cholesterol synthesis pathway are crucial for establishing β-glucan-, BCG- or oxLDL-induced trained immunity in macrophages (49). We now describe that exposure of human macrophages to antifolates (PMX or MTX) results in the acquisition of an innate tolerance state, thus demonstrating that the one-carbon metabolism is another metabolic circuit that critically mediates trained immunity.

CD14 acts both as a pattern-recognition receptor (50, 51) and is a receptor for LPS (52). In the context of RA, Lewis et al. has recently defined three distinct histopathological entities based on transcriptional data from the synovium of early treatment-naive RA patients: fibroblastic pathotype, macrophage-rich myeloid pathotype and lympho-myeloid pathotype (53). The analysis of CD14 mRNA expression in the three pathotypes (https://peac.hpc.qmul.ac.uk/) revealed that CD14 expression is higher in lymphoid pathotype (padj 5.7e-0^3^ versus myeloid pathotype, padj 6.8e-0^7^ versus fibroid pathotype), indicating that CD14 marks the lymphoid-rich pathotype with high plasma cell accumulation. Moreover, synovium CD14 expression correlated positively with disease activity (padj 0.016) (53, 54). Along the same line, sCD14 levels are increased in RA synovial fluid and serum compared to osteoarthritis patients (55, 56). sCD14 plays an important role in mediating the immune responses to LPS of CD14-negative cells such as endothelial cells and epithelial cells, and induces proinflammatory cytokines in fibroblast-like synovial cells from RA patients (57). The modulation of CD14 expression by antifolates that we now report supports the anti-inflammatory role of MTX in RA patients and leads us to suggest that MTX-treated RA patients would exhibit lower levels of sCD14 and mCD14 in myeloid cells, lower responsiveness to TLR4-dependent DAMPs, and a lower proinflammatory profile. In line with this hypothesis, we have observed that sCD14 level diminishes in serum of early rheumatoid arthritis patient’s responding to MTX treatment. Considering that MTX is the first line in RA treatment and the importance of taking advantage of the window of opportunity to achieve early remission, it would be of interest to select those patients with highest odds to be MTX responders. In this regard, a low baseline sCD14 can identify patients who are MTX non-responder, in order to try a DMARD with other mechanism of action. We are aware that there is enough overlap of sCD14 baseline levels between responder and non-responder patients. In this case it would be also useful to measure the variation in sCD14 levels that could help to better identify non-responder patients in doubtful cases.

On the other hand, it is well known that MTX withdrawal due to adverse events is more frequent than to inefficacy (58). PMX is a chemotherapeutic drug with substantial activity against lung carcinomas usually considered as refractory to classical antifolates (59). Although not used as a DMARD in RA, PMX has been shown to suppress the release of TNFα from activated T cells of RA patients, and also ameliorates experimental arthritis in a model of collagen-induced arthritis in rats, thus indicating that PMX exhibits an anti-inflammatory action both *ex vivo* and *in vivo* (10, 60). In spite of these antecedents, the anti-inflammatory efficacy of PMX has not been previously tested in the case of innate immune cells. Our results indicate that, besides a robust antifolate-mediated cytostatic effect, PMX exerts a huge reprogramming effect on human macrophages, thus opening further research of this drug to new opportunities beyond the limit of its actual clinical utility.

## ACKNOWLEDGMENTS, FUNDINGS

This work was supported by grants PI17/00037 and PI20/00316 to APK, PI18/00371 to IGA, and RIER RD16/0012/0007, RD16/0012/0011 and RD16/0012/0012 from Instituto de Salud Carlos III/FEDER to APK, MEMC and IGA, and cofinanced by European Regional Development Fund “A way to achieve Europe” (ERDF). SFR is supported by a contract from Instituto de Salud Carlos III (FI18/00109), AT-M is supported by a PhD fellowship from the Autonomous Region of Madrid (PEJD-2019-PRE/BMB-16851). APK is supported by FIBHGM.

The authors acknowledge Julia Villarejo for technical help and Dr. Angel L. Corbí, Dr. Juan Cañete and Dr. Paloma Sánchez-Mateos for helpful discussions.

## AUTHORSHIP CONTRIBUTIONS

SFR, CB, RGC, MTT, IM, ATM designed research, performed research and analyzed data; LN, AV, RGV, GJ, MEMC, IGA, designed research and analyzed data; APK conceived the study, designed research, analyzed data and wrote the paper. All authors had final approval of the version.

---

**Supplementary Figure 1.**
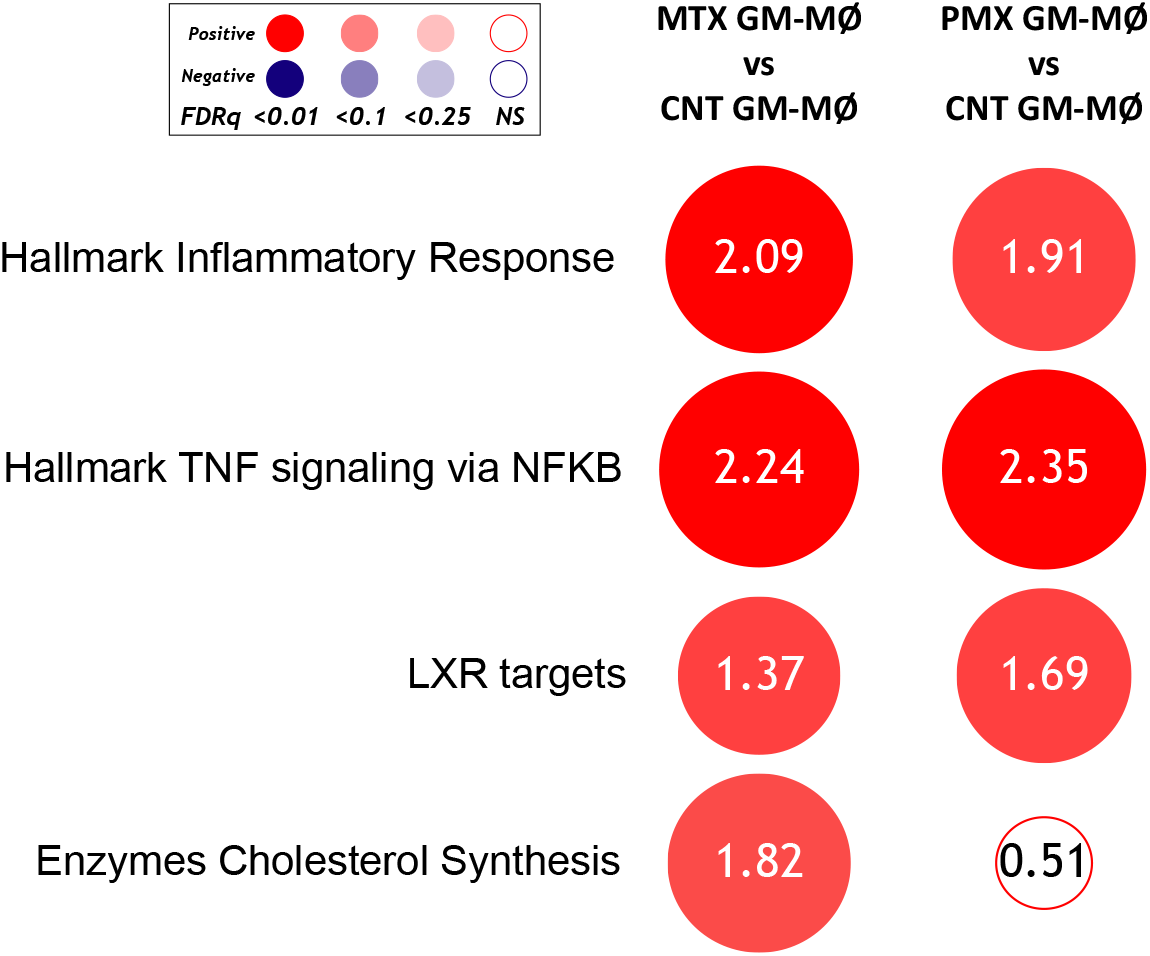
Comparison of the global transcriptional signature of the anti-folates PMX and MTX in GM-CSF-primed macrophages. Summary of GSEA with the indicated gene sets on the ranked comparison of the transcriptomes of MTX-GM-MØ vs GM-MØ, and PMX-GM-MØ vs GM-MØ. Circles area is proportional to the absolute value of the Normalized Enrichment Score (NES). The intensity of color increases with the enrichment of the gene signature (red, positive enrichment; blue, negative enrichment). False discovery rate (FDRq) is also indicated.

**Supplementary Figure 2.**
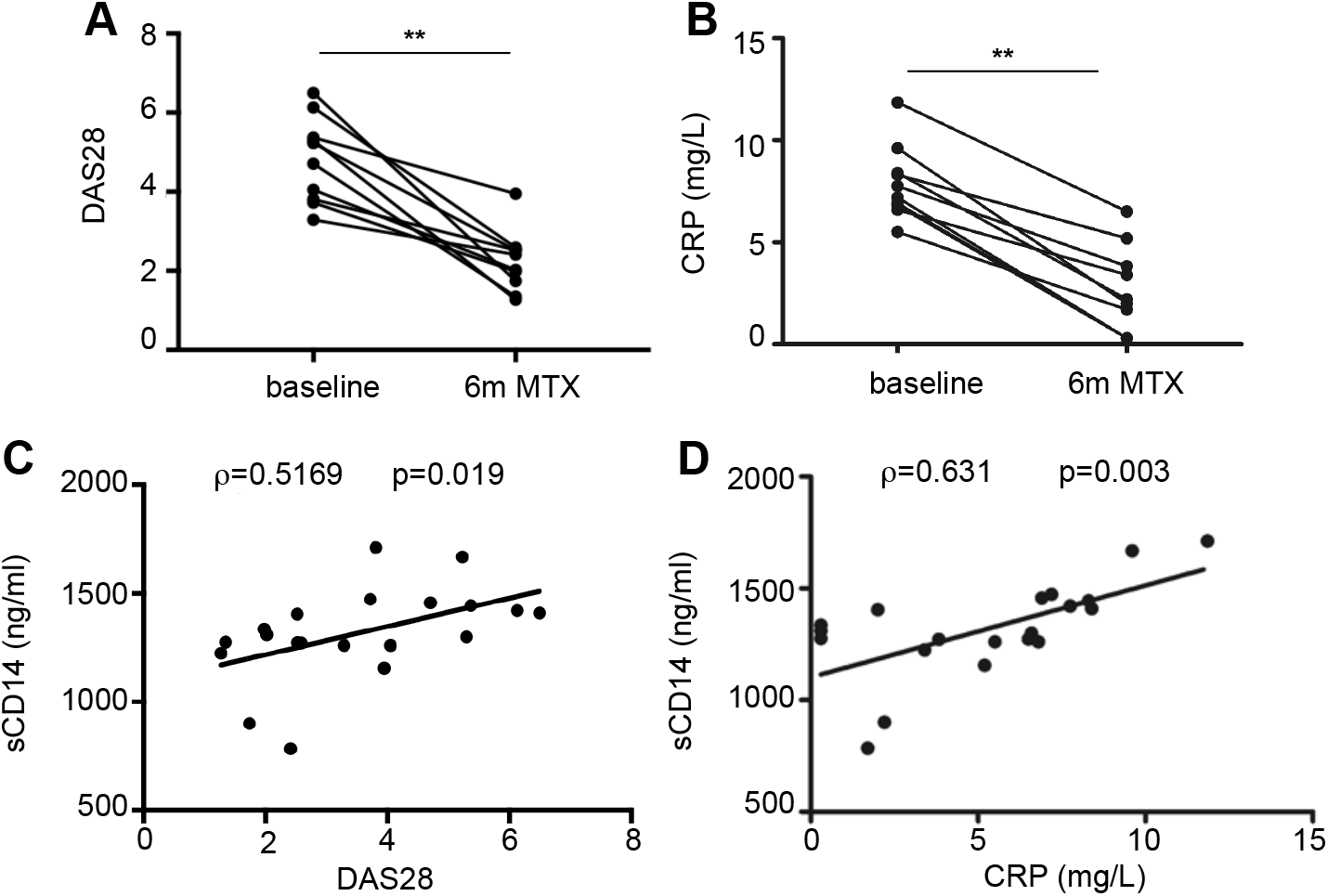
Disease activity score (DAS28) **(A)** and C-reactive protein (CRP) **(B)** from early RA patients at baseline and 6 months after MTX treatment from the discovery cohort. Correlation between sCD14 and DAS28 **(C)** sCD14 and CRP **(D)** in early RA patients (two-tailed Spearman’s correlation).

**Supplementary Table 1.** Relationship between high basal sCD14 (ng/ml) serum levels or decreased ΔsCD14 and response to Methotrexate treatment in RA patients. Multivariable logistic regression analysis to estimate with a better precision the capability of sCD14 to discriminate between MTX-responder and non-responder patients. The dependent variable was response to MTX and as independent variables we included gender, age and baseline DAS28, that are well known predictors of treatment response in RA. sCD14 or ΔsCD14 were included in the model to determine their respective OR adjusted by age, gender and baseline DAS28.

| **High basal sCD14 (ng/ml) levels** |  |  |
| --- | --- | --- |
|  | **Odds Ratio (95% CI)** | **p value** |
| **Female gender** | 0.11 (0.005 to 2.37) | 0,16 |
| **Age (years)** | 0.93 (0.85 to 1.01) | 0,1 |
| **High basal sCD14 (ng/ml) levels** | 25.49 (2.22 to 291.90) | 0,009 |
| **DAS28** | 2.11 (0.92 to 4.84) | 0,076 |
| **Decreased ΔsCD14** |  |  |
|  | **Odds Ratio (95% CI)** | **p value** |
| **Female gender** | 0.59 (0.03 to 9.58) | 0,71 |
| **Age (years)** | 0.96 (0.89 to 1.03) | 0,31 |
| **Decreased ΔsCD14** | 40.35 (3.02 to 537.38) | 0,005 |
| **DAS28** | 2.86 (1.15 to 7.12) | 0,02 |
| **Relationship between high basal sCD14 (ng/ml) serum levels or decreased ΔsCD14 and response to Methotrexate treatment in RA patients. DAS28: disease activity score estimated with 28 joint counts.** |  |  |

## REFERENCES

1. Ducker GS, and Rabinowitz JD. One-Carbon Metabolism in Health and Disease. Cell metabolism. 2017;25(1):27–42.

2. Gonen N, and Assaraf YG. Antifolates in cancer therapy: structure, activity and mechanisms of drug resistance. Drug Resist Updat. 2012;15(4):183–210.

3. Fan J, Ye J, Kamphorst JJ, Shlomi T, Thompson CB, and Rabinowitz JD. Quantitative flux analysis reveals folate-dependent NADPH production. Nature. 2014;510(7504):298–302.

4. de Gooijer CJ, Baas P, and Burgers JA. Current chemotherapy strategies in malignant pleural mesothelioma. Translational lung cancer research. 2018;7(5):574–83.

5. Hazarika M, White RM, Johnson JR, and Pazdur R. FDA drug approval summaries: pemetrexed (Alimta). The oncologist. 2004;9(5):482–8.

6. Rossi G, Alama A, Genova C, Rijavec E, Tagliamento M, Biello F, Coco S, Dal Bello MG, Boccardo S, and Grossi F. The evolving role of pemetrexed disodium for the treatment of non-small cell lung cancer. Expert opinion on pharmacotherapy. 2018;19(17):1969–76.

7. Qiu A, Jansen M, Sakaris A, Min SH, Chattopadhyay S, Tsai E, Sandoval C, Zhao R, Akabas MH, and Goldman ID. Identification of an intestinal folate transporter and the molecular basis for hereditary folate malabsorption. Cell. 2006;127(5):917–28.

8. Shih C, Chen VJ, Gossett LS, Gates SB, MacKellar WC, Habeck LL, Shackelford KA, Mendelsohn LG, Soose DJ, Patel VF, et al. LY231514, a pyrrolo[2,3-d]pyrimidine-based antifolate that inhibits multiple folate-requiring enzymes. Cancer Res. 1997;57(6):1116–23.

9. Smolen JS, Landewe R, Bijlsma J, Burmester G, Chatzidionysiou K, Dougados M, Nam J, Ramiro S, Voshaar M, van Vollenhoven R, et al. EULAR recommendations for the management of rheumatoid arthritis with synthetic and biological disease-modifying antirheumatic drugs: 2016 update. Ann Rheum Dis. 2017;76(6):960–77.

10. Karatas A, Koca SS, Ozgen M, Dagli AF, Erman F, Sahin N, Sahin K, and Isik A. Pemetrexed ameliorates experimental arthritis in rats. Inflammation. 2015;38(1):9–15.

11. Netea MG, Quintin J, and van der Meer JW. Trained immunity: a memory for innate host defense. Cell Host Microbe. 2011;9(5):355–61.

12. Netea MG, Joosten LA, Latz E, Mills KH, Natoli G, Stunnenberg HG, O’Neill LA, and Xavier RJ. Trained immunity: A program of innate immune memory in health and disease. Science. 2016;352(6284):aaf1098.

13. Netea MG, Dominguez-Andres J, Barreiro LB, Chavakis T, Divangahi M, Fuchs E, Joosten LAB, van der Meer JWM, Mhlanga MM, Mulder WJM, et al. Defining trained immunity and its role in health and disease. Nature reviews Immunology. 2020.

14. Song WM, and Colonna M. Immune Training Unlocks Innate Potential. Cell. 2018;172(1-2):3–5.

15. Choy EH, and Panayi GS. Cytokine pathways and joint inflammation in rheumatoid arthritis. N Engl J Med. 2001;344(12):907–16.

16. Scott DL, Wolfe F, and Huizinga TW. Rheumatoid arthritis. Lancet. 2010;376(9746):1094–108.

17. McInnes IB, and Schett G. The pathogenesis of rheumatoid arthritis. N Engl J Med. 2011;365(23):2205–19.

18. Soler Palacios B, Estrada-Capetillo L, Izquierdo E, Criado G, Nieto C, Municio C, Gonzalez-Alvaro I, Sanchez-Mateos P, Pablos JL, Corbi AL, et al. Macrophages from the synovium of active rheumatoid arthritis exhibit an activin A-dependent pro-inflammatory profile. J Pathol. 2015;235(3):515–26.

19. Municio C, Soler Palacios B, Estrada-Capetillo L, Benguria A, Dopazo A, Garcia-Lorenzo E, Fernandez-Arroyo S, Joven J, Miranda-Carus ME, Gonzalez-Alvaro I, et al. Methotrexate selectively targets human proinflammatory macrophages through a thymidylate synthase/p53 axis. Ann Rheum Dis. 2016;75(12):2157–65.

20. Municio C, Dominguez-Soto A, Fuentelsaz-Romero S, Lamana A, Montes N, Cuevas VD, Campos RG, Pablos JL, Gonzalez-Alvaro I, and Puig-Kroger A. Methotrexate limits inflammation through an A20-dependent cross-tolerance mechanism. Ann Rheum Dis. 2018;77(5):752–9.

21. Visser K, Katchamart W, Loza E, Martinez-Lopez JA, Salliot C, Trudeau J, Bombardier C, Carmona L, van der Heijde D, Bijlsma JW, et al. Multinational evidence-based recommendations for the use of methotrexate in rheumatic disorders with a focus on rheumatoid arthritis: integrating systematic literature research and expert opinion of a broad international panel of rheumatologists in the 3E Initiative. Ann Rheum Dis. 2009;68(7):1086–93.

22. Aletaha D, Neogi T, Silman AJ, Funovits J, Felson DT, Bingham CO, 3rd, Birnbaum NS, Burmester GR, Bykerk VP, Cohen MD, et al. 2010 rheumatoid arthritis classification criteria: an American College of Rheumatology/European League Against Rheumatism collaborative initiative. Ann Rheum Dis. 2010;69(9):1580–8.

23. Prevoo ML, van’t Hof MA, Kuper HH, van Leeuwen MA, van de Putte LB, and van Riel PL. Modified disease activity scores that include twenty-eight-joint counts. Development and validation in a prospective longitudinal study of patients with rheumatoid arthritis. Arthritis Rheum. 1995;38(1):44–8.

24. Tate J, and Ward G. Interferences in immunoassay. The Clinical biochemist Reviews. 2004;25(2):105–20.

25. Gonzalez-Alvaro I, Ortiz AM, Alvaro-Gracia JM, Castaneda S, Diaz-Sanchez B, Carvajal I, Garcia-Vadillo JA, Humbria A, Lopez-Bote JP, Patino E, et al. Interleukin 15 levels in serum may predict a severe disease course in patients with early arthritis. PLoS One. 2011;6(12):e29492.

26. Langmead B, and Salzberg SL. Fast gapped-read alignment with Bowtie 2. Nat Methods. 2012;9(4):357–9.

27. Li B, and Dewey CN. RSEM: accurate transcript quantification from RNA-Seq data with or without a reference genome. BMC Bioinformatics. 2011;12(323.

28. Subramanian A, Tamayo P, Mootha VK, Mukherjee S, Ebert BL, Gillette MA, Paulovich A, Pomeroy SL, Golub TR, Lander ES, et al. Gene set enrichment analysis: a knowledge-based approach for interpreting genome-wide expression profiles. Proc Natl Acad Sci U S A. 2005;102(43):15545–50.

29. Fischer M. Census and evaluation of p53 target genes. Oncogene. 2017;36(28):3943–56.

30. Edgar R, Domrachev M, and Lash AE. Gene Expression Omnibus: NCBI gene expression and hybridization array data repository. Nucleic acids research. 2002;30(1):207–10.

31. Matherly LH, Hou Z, and Deng Y. Human reduced folate carrier: translation of basic biology to cancer etiology and therapy. Cancer metastasis reviews. 2007;26(1):111–28.

32. Rackov G, Hernandez-Jimenez E, Shokri R, Carmona-Rodriguez L, Manes S, Alvarez-Mon M, Lopez-Collazo E, Martinez AC, and Balomenos D. p21 mediates macrophage reprogramming through regulation of p50-p50 NF-kappaB and IFN-beta. J Clin Invest. 2016;126(8):3089–103.

33. Biswas SK, and Lopez-Collazo E. Endotoxin tolerance: new mechanisms, molecules and clinical significance. Trends Immunol. 2009;30(10):475–87.

34. Park SH, Park-Min KH, Chen J, Hu X, and Ivashkiv LB. Tumor necrosis factor induces GSK3 kinase-mediated cross-tolerance to endotoxin in macrophages. Nature immunology. 2011;12(7):607–15.

35. Yang TY, Chang GC, Chen KC, Hung HW, Hsu KH, Sheu GT, and Hsu SL. Sustained activation of ERK and Cdk2/cyclin-A signaling pathway by pemetrexed leading to S-phase arrest and apoptosis in human non-small cell lung cancer A549 cells. European journal of pharmacology. 2011;663(1-3):17–26.

36. Hacker H, Redecke V, Blagoev B, Kratchmarova I, Hsu LC, Wang GG, Kamps MP, Raz E, Wagner H, Hacker G, et al. Specificity in Toll-like receptor signalling through distinct effector functions of TRAF3 and TRAF6. Nature. 2006;439(7073):204–7.

37. Nomura F, Akashi S, Sakao Y, Sato S, Kawai T, Matsumoto M, Nakanishi K, Kimoto M, Miyake K, Takeda K, et al. Cutting edge: endotoxin tolerance in mouse peritoneal macrophages correlates with down-regulation of surface toll-like receptor 4 expression. J Immunol. 2000;164(7):3476–9.

38. Murphy MB, Xiong Y, Pattabiraman G, Manavalan TT, Qiu F, and Medvedev AE. Pellino-3 promotes endotoxin tolerance and acts as a negative regulator of TLR2 and TLR4 signaling. J Leukoc Biol. 2015;98(6):963–74.

39. Van Overmeire E, Stijlemans B, Heymann F, Keirsse J, Morias Y, Elkrim Y, Brys L, Abels C, Lahmar Q, Ergen C, et al. M-CSF and GM-CSF Receptor Signaling Differentially Regulate Monocyte Maturation and Macrophage Polarization in the Tumor Microenvironment. Cancer Res. 2016;76(1):35–42.

40. Hamilton JA. Colony-stimulating factors in inflammation and autoimmunity. Nature reviews Immunology. 2008;8(7):533–44.

41. Brown PM, Pratt AG, and Isaacs JD. Mechanism of action of methotrexate in rheumatoid arthritis, and the search for biomarkers. Nature reviews Rheumatology. 2016;12(12):731–42.

42. Zanoni I, Ostuni R, Marek LR, Barresi S, Barbalat R, Barton GM, Granucci F, and Kagan JC. CD14 controls the LPS-induced endocytosis of Toll-like receptor 4. Cell. 2011;147(4):868–80.

43. Perera PY, Mayadas TN, Takeuchi O, Akira S, Zaks-Zilberman M, Goyert SM, and Vogel SN. CD11b/CD18 acts in concert with CD14 and Toll-like receptor (TLR) 4 to elicit full lipopolysaccharide and taxol-inducible gene expression. J Immunol. 2001;166(1):574–81.

44. Palsson-McDermott EM, and O’Neill LAJ. Targeting immunometabolism as an anti-inflammatory strategy. Cell research. 2020;30(4):300–14.

45. Dominguez-Andres J, Joosten LA, and Netea MG. Induction of innate immune memory: the role of cellular metabolism. Current opinion in immunology. 2019;56(10–6.

46. Cheng SC, Quintin J, Cramer RA, Shepardson KM, Saeed S, Kumar V, Giamarellos-Bourboulis EJ, Martens JH, Rao NA, Aghajanirefah A, et al. mTOR- and HIF-1alpha-mediated aerobic glycolysis as metabolic basis for trained immunity. Science. 2014;345(6204):1250684.

47. Arts RJW, Carvalho A, La Rocca C, Palma C, Rodrigues F, Silvestre R, Kleinnijenhuis J, Lachmandas E, Goncalves LG, Belinha A, et al. Immunometabolic Pathways in BCG-Induced Trained Immunity. Cell reports. 2016;17(10):2562–71.

48. Dominguez-Andres J, Novakovic B, Li Y, Scicluna BP, Gresnigt MS, Arts RJW, Oosting M, Moorlag S, Groh LA, Zwaag J, et al. The Itaconate Pathway Is a Central Regulatory Node Linking Innate Immune Tolerance and Trained Immunity. Cell metabolism. 2019;29(1):211–20 e5.

49. Bekkering S, Arts RJW, Novakovic B, Kourtzelis I, van der Heijden C, Li Y, Popa CD, Ter Horst R, van Tuijl J, Netea-Maier RT, et al. Metabolic Induction of Trained Immunity through the Mevalonate Pathway. Cell. 2018;172(1-2):135–46 e9.

50. Zanoni I, and Granucci F. Role of CD14 in host protection against infections and in metabolism regulation. Frontiers in cellular and infection microbiology. 2013;3(32.

51. Leveque M, Simonin-Le Jeune K, Jouneau S, Moulis S, Desrues B, Belleguic C, Brinchault G, Le Trionnaire S, Gangneux JP, Dimanche-Boitrel MT, et al. Soluble CD14 acts as a DAMP in human macrophages: origin and involvement in inflammatory cytokine/chemokine production. FASEB journal : official publication of the Federation of American Societies for Experimental Biology. 2017;31(5):1891–902.

52. Zanoni I, Tan Y, Di Gioia M, Springstead JR, and Kagan JC. By Capturing Inflammatory Lipids Released from Dying Cells, the Receptor CD14 Induces Inflammasome-Dependent Phagocyte Hyperactivation. Immunity. 2017;47(4):697–709 e3.

53. Lewis MJ, Barnes MR, Blighe K, Goldmann K, Rana S, Hackney JA, Ramamoorthi N, John CR, Watson DS, Kummerfeld SK, et al. Molecular Portraits of Early Rheumatoid Arthritis Identify Clinical and Treatment Response Phenotypes. Cell reports. 2019;28(9):2455–70 e5.

54. Humby F, Lewis M, Ramamoorthi N, Hackney JA, Barnes MR, Bombardieri M, Setiadi AF, Kelly S, Bene F, DiCicco M, et al. Synovial cellular and molecular signatures stratify clinical response to csDMARD therapy and predict radiographic progression in early rheumatoid arthritis patients. Ann Rheum Dis. 2019;78(6):761–72.

55. Smiljanovic B, Radzikowska A, Kuca-Warnawin E, Kurowska W, Grun JR, Stuhlmuller B, Bonin M, Schulte-Wrede U, Sorensen T, Kyogoku C, et al. Monocyte alterations in rheumatoid arthritis are dominated by preterm release from bone marrow and prominent triggering in the joint. Ann Rheum Dis. 2018;77(2):300–8.

56. Smiljanovic B, Grutzkau A, Sorensen T, Grun JR, Vogl T, Bonin M, Schendel P, Stuhlmuller B, Claussnitzer A, Hermann S, et al. Synovial tissue transcriptomes of long-standing rheumatoid arthritis are dominated by activated macrophages that reflect microbial stimulation. Sci Rep. 2020;10(1):7907.

57. Ichise Y, Saegusa J, Tanaka-Natsui S, Naka I, Hayashi S, Kuroda R, and Morinobu A. Soluble CD14 Induces Pro-inflammatory Cytokines in Rheumatoid Arthritis Fibroblast-Like Synovial Cells via Toll-Like Receptor 4. Cells. 2020;9(7).

58. Lopez-Olivo MA, Siddhanamatha HR, Shea B, Tugwell P, Wells GA, and Suarez-Almazor ME. Methotrexate for treating rheumatoid arthritis. The Cochrane database of systematic reviews. 20146):CD000957.

59. Scagliotti GV, Parikh P, von Pawel J, Biesma B, Vansteenkiste J, Manegold C, Serwatowski P, Gatzemeier U, Digumarti R, Zukin M, et al. Phase III study comparing cisplatin plus gemcitabine with cisplatin plus pemetrexed in chemotherapy-naive patients with advanced-stage non-small-cell lung cancer. J Clin Oncol. 2008;26(21):3543–51.

60. van der Heijden JW, Assaraf YG, Gerards AH, Oerlemans R, Lems WF, Scheper RJ, Dijkmans BA, and Jansen G. Methotrexate analogues display enhanced inhibition of TNF-alpha production in whole blood from RA patients. Scandinavian journal of rheumatology. 2014;43(1):9–16.

